# Hybrid-Lambda: simulation of multiple merger and Kingman gene genealogies in species networks and species trees

**DOI:** 10.1101/023465

**Authors:** Sha Zhu, James H. Degnan, Sharyn J. Goldstien, Bjarki Eldon

## Abstract

**Background:** There has been increasing interest in coalescent models which admit multiple mergers of ancestral lineages; and to model hybridization and coalescence simultaneously.

**Results:** `Hybrid-Lambda` is a software package that simulates gene genealogies under multiple merger and Kingman’s coalescent processes within species networks or species trees. `Hybrid-Lambda` allows different coalescent processes to be specified for different populations, and allows for time to be converted between generations and coalescent units, by specifying a population size for each population. In addition, `Hybrid-Lambda` can generate simulated datasets, assuming the infinitely many sites mutation model, and compute the *F*_*ST*_ statistic. As an illustration, we apply `Hybrid-Lambda` to infer the time of subdivision of certain marine invertebrates under different coalescent processes.

**Conclusions:** `Hybrid-Lambda` makes it possible to investigate biogeographic concordance among high fecundity species exhibiting skewed offspring distribution.

## Background

Species trees describe ancestral relations among species. Gene genealogies describe the random ancestral relations of alleles sampled within species. Species trees are often assumed to be bifurcating [6], and gene genealogies to follow the Kingman coalescent [23, 28] in allowing at most two lineages to coalesce at a time.

Recently, there has been increasing interest in coalescent models which admit multiple mergers of ancestral lineages [1, 2, 9, 12, 37, 39, 40] and to model hybridization and coalescence simultaneously [3, 25, 26, 29, 46]. For high fecundity species exhibiting sweepstake-like reproduction, such as oysters and other marine organisms [1, 4, 9, 11, 17, 18, 39], the Kingman coalescent may not be appropriate, as it is based on low offspring number population models (see recent reviews by Hedgecock and Pudovkin [19] and Tellier and Lemaire [42]). Thus, we consider Λ coalescents [8, 36, 37] derived from sweepstake-like reproduction models, and allow *more* than two lineages to coalesce at a time. We introduce the software `Hybrid-Lambda` for simulating gene trees under two models of Λ-coalescents within rooted species trees and rooted species networks. Our program differs from existing software which also allows multiple mergers, such as SIMCOAL 2.0 [30] — which allows multiple mergers in gene trees due to small population sizes under the Wright-Fisher model — in that we apply coalescent processes that are obtained from population models explicitly modelling skewed offspring distributions, as opposed to bottlenecks.

Species trees may also fail to be bifurcating due to either polytomies or hybridization events. The simulation of gene genealogies within a species network which admits hybridization is another application of `Hybrid-Lambda`. The package `ms` [24] can also simulate gene genealogies within species networks under Kingman’s coalescent. However the input of `ms` is difficult to automate when the network is sophisticated or generated from other software. Other simulation studies using species networks have either used a small number of network topologies coded individually (for example, in `phylonet` [43, 45, 46]) or have assumed that gene trees have evolved on species trees embedded within the species network [22, 29, 32]. `Hybrid-Lambda` will help to automate simulation studies of hybridization by allowing for a large number of species network topologies and allowing gene trees to evolve directly within the network. `Hybrid-Lambda` can simulate both Kingman and Λ-coalescent processes within species networks. A comparison of features of several software packages that output gene genealogies under coalescent models is given in Table 1.

**Table 1.** **Comparison of software programs simulating gene trees in species trees and networks.** *Migration* **refers to modeling post-speciation gene flow.**

| | gene trees<br>in networks | Λ-coalescent | small pop.<br>mult. merger | infinite sites<br>model | recombination | migration |
| --- | --- | --- | --- | --- | --- | --- |
| COAL[6] | no | no | no | no | no | no |
| CoaSim[33] | no | no | no | yes | yes | yes |
| fastsimcoal2[15] | yes | no | yes | no | yes | yes |
| Genome[31] | yes | no | no | yes | yes | yes |
| GUMS[20] | no | no | no | no | no | yes |
| ms[24] | yes | no | no | yes | yes | yes |
| msHOT[21] | yes | no | no | yes | yes | yes |
| msms[14] | yes | no | no | yes | yes | yes |
| scrm[41] | yes | no | no | yes | yes | yes |
| simcoal2[27] | yes | no | yes | no | yes | yes |
| hybrid-Lambda | yes | yes | no | yes | no | no |

## Implementation

The program input file for `Hybrid-Lambda` is a character string that describes relationships between species. Standard Newick format [34] is used for the input of species trees and the output of gene trees, whose interior nodes are not labelled. An extended Newick formatted string [5, 25] labels all internal nodes, and is used for the input of species networks (see Fig. 1).

**Figure 1.**
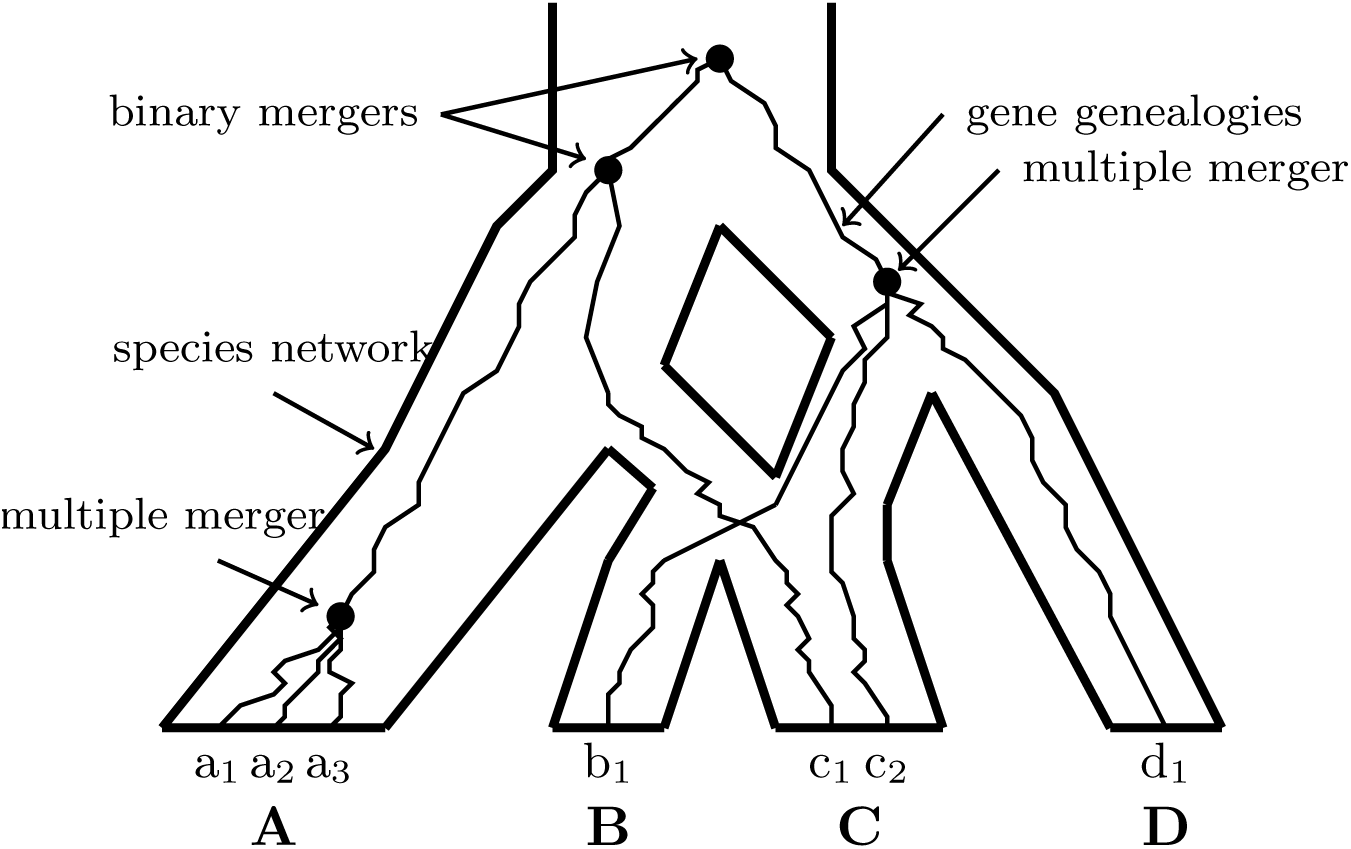
Demonstration of a multiple merger genealogy within a specis network. A multiple merger gene genealogy with topology (((a_1_,a_2_,a_3_),c_1_),(b_1_,c_2_,d_1_)), of which the coalescence events pointed to by arrows labelled “multiple merger” indicate coalescence of 3 lines, simulated in a species network with topology `((((B,C)s1)h1#H1,A)s2,(h1#H1,D)s3)r, where H1` is the probability that a lineage has its ancestry from its left parental population.

### Parameters

`Hybrid-Lambda` can use multiple lineages sampled from each species and simulate Kingman or multiple merger (Λ)-coalescent processes within a given species network. In addition, separate coalescent processes can be specified on different branches of the species network. The coalescent is a continuous-time Markov process, in which times between coalescent events are independent exponential random variables with different rates. The rates are determined by a so-called coalescent parameter that can be input via command line, or a(n) (extended) Newick formatted string with specific coalescent parameters as branch lengths. By default, the Kingman coalescent is used, for which two of *b* active lineages coalesce at rate 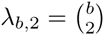. One can choose between two different examples of a Λ-coalescent, whose parameters have clear biological interpretation. While we cannot hope to cover the huge class of Lambda-coalescents, our two examples are the ones that have been most studied in the literature [2, 7, 13]. If the coalescent parameter is between 0 and 1, then we use *ψ* for the coalescent parameter, and the rate *λ*_*b,k*_ at which *k* out of *b* (2 *≤ k ≤ b*) active ancestral lineages merge is

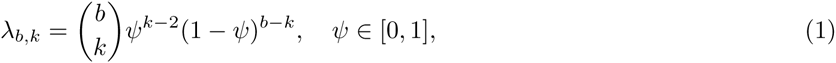

[9]. If the coalescent parameter is between 1 and 2, then we use *α* for the coalescent parameter, and the rate of *k*-mergers (2 *≤ k ≤ b*) is

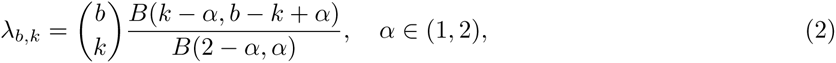

where *B*(*·, ·*) is the beta function [40].

`Hybrid-Lambda` assumes by default that the input network (tree) branch lengths are in coalescent units. However, this is not essential. Coalescent units can be converted through an alternative input file with numbers of generations as branch lengths, which are then divided by their corresponding effective population sizes. By default, effective population sizes on all branches are assumed to be equal and unchanged. Users can change this parameter using the command line, or using a(n) (extended) Newick formatted string to specify population sizes on all branches through another input file.

The simulation requires ultrametric species networks, i.e. equal lengths of all paths from tip to root. `Hybrid-Lambda` checks the distances in coalescent units between the root and all tip nodes and prints out warning messages if the ultrametric assumption is violated.

## Results and Discussion

`Hybrid-Lambda` outputs simulated gene trees in three different files: one contains gene trees with branch lengths in coalescent units, another uses the number of generations as branch lengths, and the third uses the number of expected mutations as branch lengths.

Besides outputting gene tree files, `Hybrid-Lambda` also provides several functions for analysis purposes:

- user-defined random seed for simulation,
- output simulated data in 0/1 format assuming the infinitely many sites mutation model,
- a frequency table of gene tree topologies,
- a figure of the species network or tree (this function only works when LATEX or `dot` is installed) (Fig. 2),
- the expected *F*_*ST*_ value for a split model between two populations,
- when gene trees are simulated from two populations, the software `Hybrid-Lambda` can generate a table of relative frequencies of reciprocal monophyly, paraphyly, and polyphyly.

**Figure 2.**
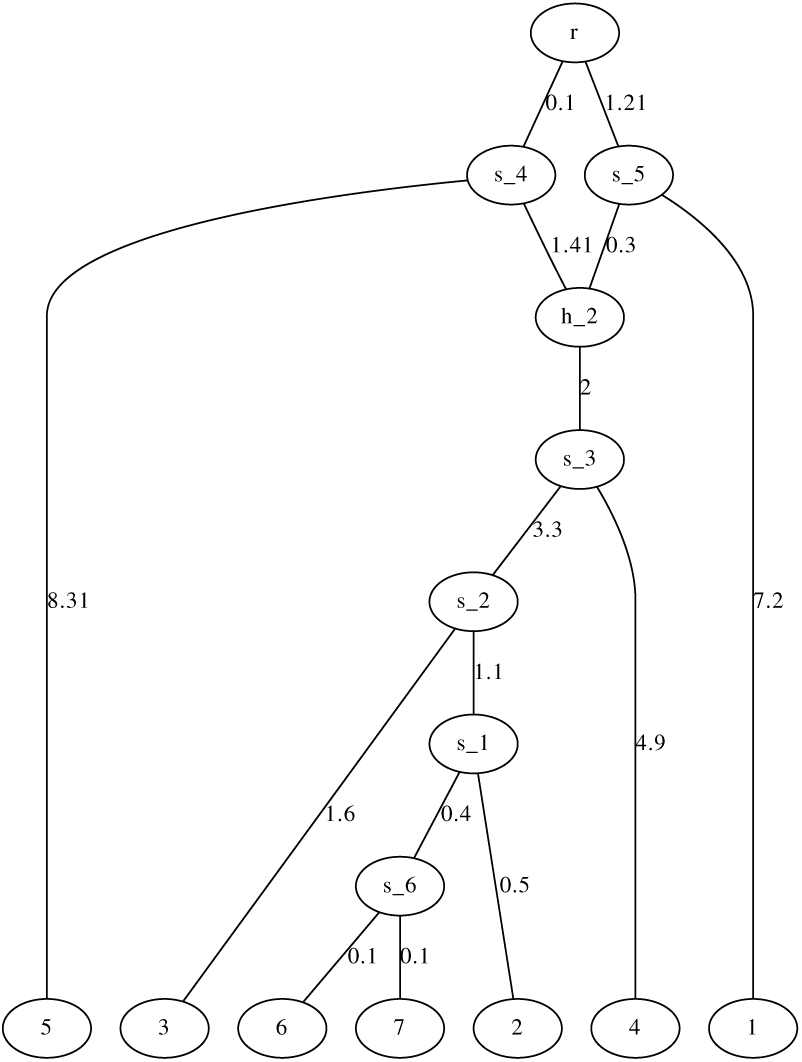
Demonstration of a network figure generated by Hybrid-Lambda. The network is automatically generated by Hybrid-Lambda as dot and. pdf files from the extended newick string “`(((((((6:.1,7:.1)s 6:.4,2:.5)s 1:1.1,3:1.6)s 2:3.3, 4:4.9)s 3:2)h 2#.5:1.41,5:8.31)s 4:0.1,(1:7.2,h 2#.5:.3)s 5:1.21)r;`”

### Simulation Example

We give a simulation example showing the impact of the particular coalescent model on estimating the divergence time for two populations. Results can be confirmed using analytic approximations to *F*_*ST*_. This is shown in the Appendix along with example code for using `Hybrid-Lambda` for this example.

Eldon and Wakeley [10] showed that population subdivision can be observed in genetic data despite high migration between populations. One of the most widely used measures of population differentiation is the *F*_*ST*_ statistic. The relationship between *F*_*ST*_ and biogeography depends on the underlying coalescent process, which might be especially important for the interpretation of divergence and demographic history of many marine species. Here we used `Hybrid-Lambda` to simulate divergence between two populations based on different Λ-coalescents, as well as the standard Kingman coalescent. Mutations were simulated in `Hybrid-Lambda` under the infinite-sites model. The summary statistic *F*_*ST*_ was estimated for these data and was used to compare *F*_*ST*_ estimated from mtDNA from five species of marine invertebrates. These species were used in previous studies to test the hypothesis that contemporary oceanic conditions are creating subdivisions between the North Island and South Island reef populations of New Zealand [16, 35, 44]. These studies represent some of the earliest mitochondrial studies on the marine disjunction between the North and South Islands of New Zealand.

The *F*_*ST*_ statistic between North Island and South Island populations reported for these species ranges from approximately 0.07 to 0.8 (Fig. 3). *Cellana ornata* displays a very strong split, which was estimated to have occurred around 0.2–0.3 million years ago based on published estimates of divergence rates and reciprocal monophyly displayed in the data set. This result may be supported by our simulations using the Kingman coalescent. However, when multiple mergers and a higher fraction of replacement by a single parent is allowed to occur then our simulations support much younger splits between the populations *∼*9,000 generations or *∼*48,000 generations ago (Fig. 3). Similarly, the strong split observed for *Coscinasterias muricata* could be placed anywhere from *∼*9000 to 45,000 generations ago depending on the degree to which multiple mergers are allowed to occur. While the range for *Patiriella regularis*, *Cellana radians* and *C. flava* is much smaller, it is still not clear cut as to whether divergence would be observed under different coalescent models. Here we used *ψ* = 0.01 and *ψ* = 0.23, and *α* = 1.5 and *α* = 1.9, with larger values of *ψ* and smaller values of *α* corresponding to higher probabilities of multiple mergers. Our choice of parameer values corresponds to the estimated values obtained for mtDNA of oysters and Atlantic cod. An estimate for Pacific oysters based on mitochondrial DNA for *ψ* was 0.075 [9]. The results for our choice of parameter values suggest that our conclusions about a much earlier split of the populations than previously estimated are robust with regard to parameter choice. A recent study of Atlantic cod [2] estimated *ψ* between 0.07 and 0.23 for nuclear genes and near 0.01 for mitochondrial genes. The same study estimated *α* to be 1.0 and 1.28 for nuclear genes and between 1.53 and 2.0 for mitochondrial genes.

**Figure 3.**
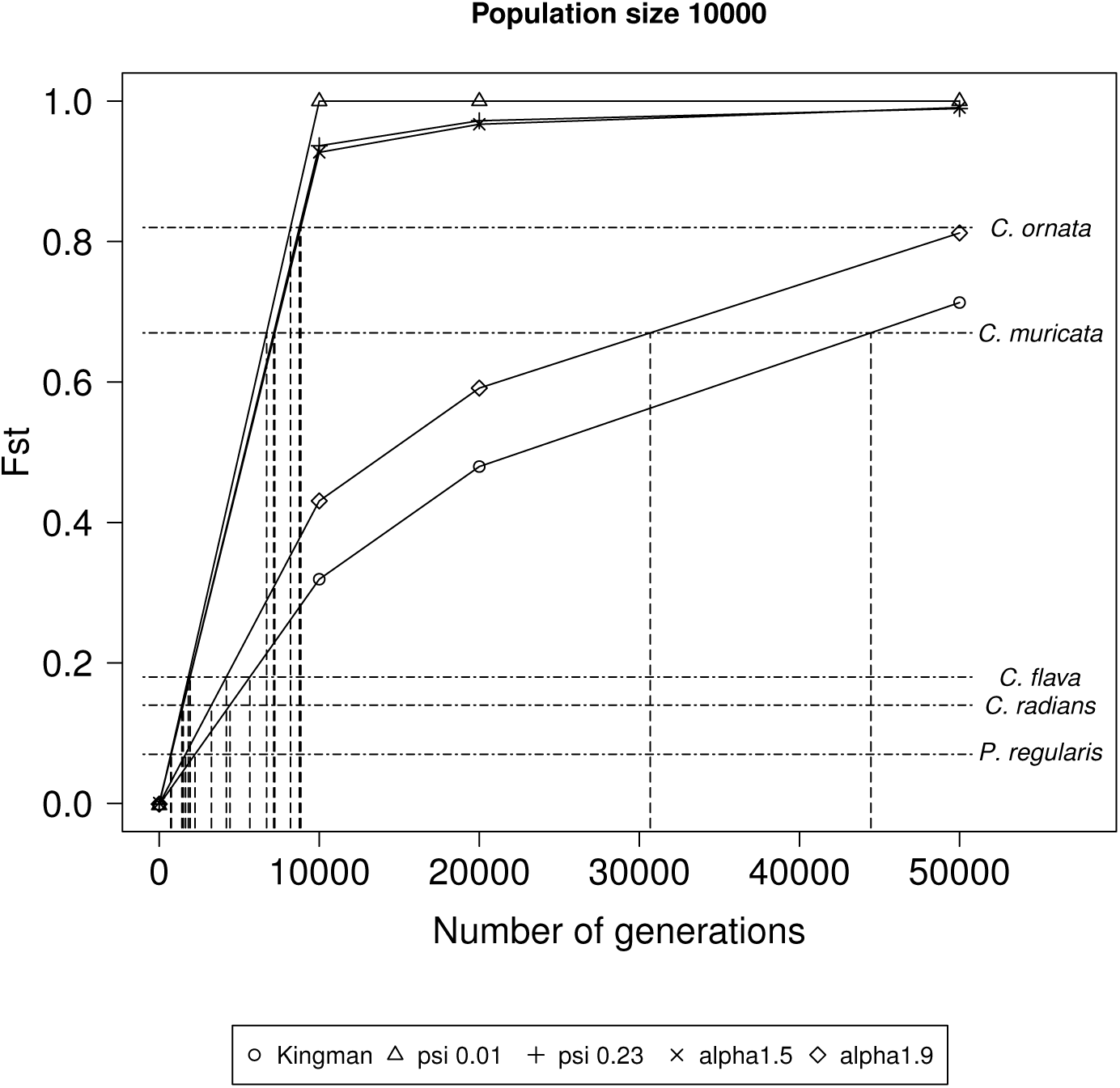
Estimated *F*_*ST*_ from simulation. The estimated *F*_*ST*_ from two populations simulated to have diverged over 0, 10, 20, and 50 thousand generations, as a function of the underlying coalescent process. Dashed lines show the relationship between the *F*_*ST*_ value estimated from mtDNA data and the estimated number of generations since divergence, for the different coalescent processes for the five marine invertebrate species, *Cellana ornata*, *C. radians*, *C. flava* (Goldstien et al. 2006), *Coscinasterias muricata* (Perrin et al. 2004), and *Patiriella regularis* (Waters and Roy 2005).

**Figure 4.**
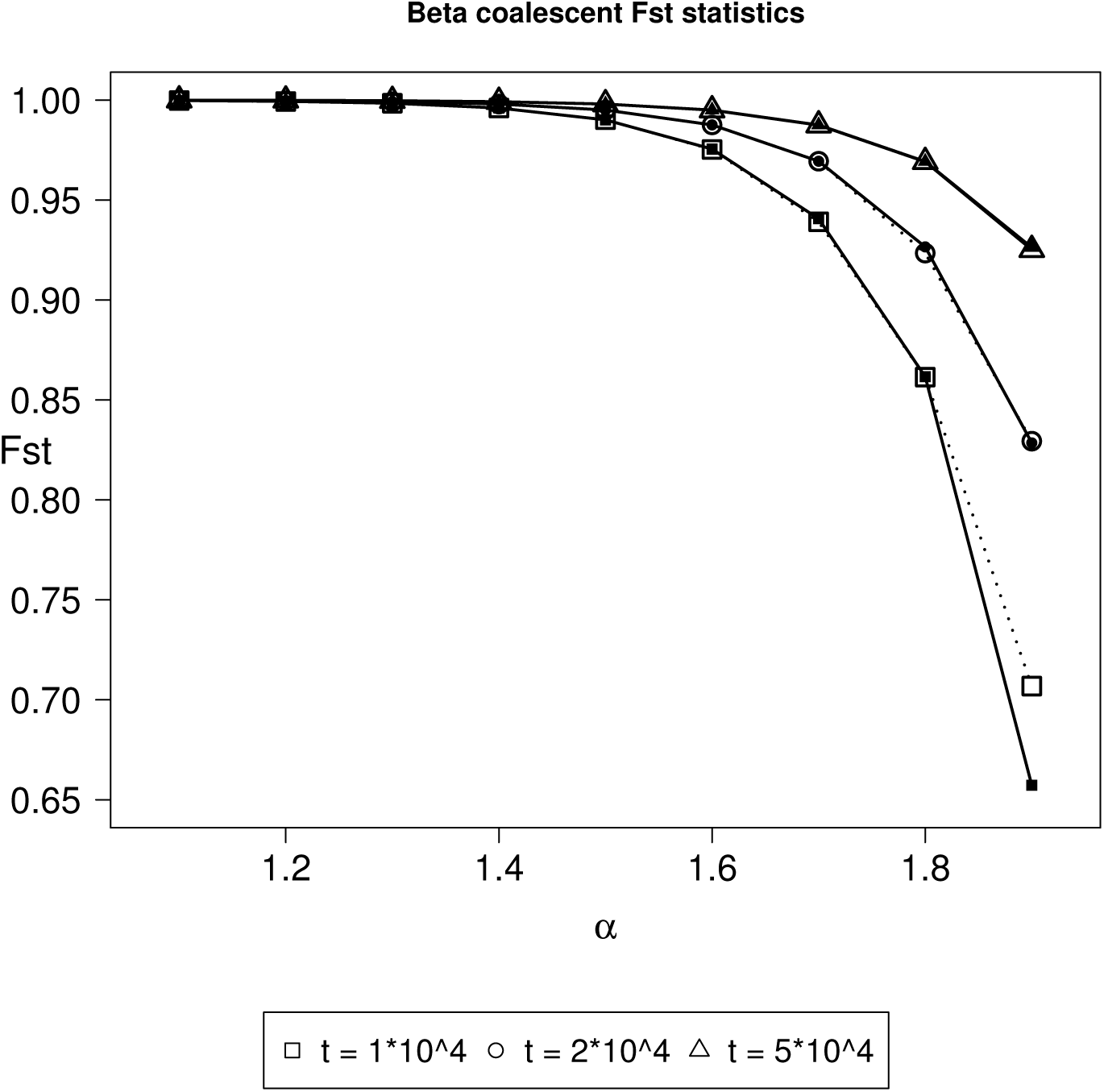
Comparison of estimated *F*_*ST*_ values from simulation and analytical predictions. Values of *F*_*ST*_ as a function of the parameter *α* for 1 *< α <* 2 for different numbers of generations of separation for two populations. Simulations (dotted lines) are based on 1 individual from each of two populations separated by *t* generations with 103 replicates and *α ∈* {1.1, 1.2, *…,* 1.9}. Analytical predictions (solid lines) of *F*_*ST*_ were calculated using (6).

## Conclusions

The implications for using alternative coalescent models are far reaching. Many marine organisms reproduce through broadcast spawning of thousands to millions of gametes, and while the expected survival of these offspring is low, there is the potential for a small subset of the adults to have a greater contribution to the next generation than assumed by the Kingman coalescent. `Hybrid-Lambda` makes it possible to investigate the effect of high fecundity on biogeographic concordance among species that exhibit high fecundity and high offspring mortality, including in complex demagraphic scenarios that allow hybridization.

## Availability and requirements

`Hybrid-Lambda` can be downloaded from http://hybridlambda.github.io/. The program is written in `C++` (requires compilers that support C++11 standard to build), and released under the GNU General Public License (GPL) version 3 or later. Users can modify and make new distributions under the terms of this license. For full details of this license, visit http://www.gnu.org/licenses/. We have used travis continuous integration to test compiling the program on Linux and Mac OS. An API in `R` [38] is currently under development.

## Author’s contributions

SZ was responsible for the software development. JD and BE supervised the project. BE derived all the *F*_*ST*_ calculations in the appendix. SG provided the simulation results and time estimates in Figure 3. All the authors have contributed to the manuscript writing.

## Acknowledgements

This work was supported by New Zealand Marsden Fund (SZ and JD), EPSRC grant EP/G052026/1 and DFG grant BL 1105/3-1 through the SPP Priority Programme 1590 “Probabilistic Structures in Evolution” (BE). This work was partly conducted while JD was a Sabbatical Fellow at the National Institute for Mathematical and Biological Synthesis, an Institute sponsored by the National Science Foundation, the U.S. Department of Homeland Security, and the U.S. Department of Agriculture through NSF Award #EF-0832858, with additional support from The University of Tennessee, Knoxville.

## Appendix *F*_*ST*_ calculations

Here we show analytic calculations that can be used to obtain expressions for *F*_*ST*_ when mutation rates are low. The effect of *α* on *F*_*ST*_ for fixed generation times is shown in Figure 4.

Assume two populations *A* and *B* have been isolated until time *τ* in the past as measured from the present. Assume also that the same coalescent process is operating in populations *A* and *B*. Let *T*_*w*_ denote the time until coalescence for two lines when drawn from the same population, and *T*_*b*_ when drawn from different populations. Let *λ*_*A*_ denote the coalescence rate for two lines in population *A*, and *λ*_*AB*_ for the common ancestral population *AB*. For the Beta(2 *- α, α*)-coalescent, *λ*_*A*_ = 1, for the point-mass process *λ*_*A*_ = *ψ*^2^. One now obtains

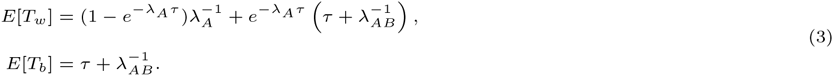

Slatkin (1991) obtained the approximation, where *μ* is the per generation mutation rate,

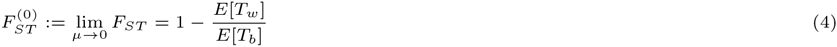

Thus, using (3) gives

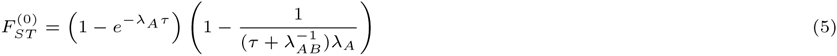

The result (5) seems to make sense, since 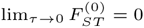 and 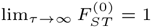. By way of example, if all populations exhibit a Beta(2 *- α, α*)-coalescent, *λ*_*A*_ = *λ*_*AB*_ = 1, and

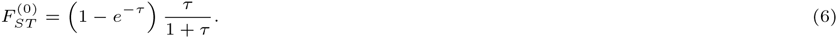

However, deciding the timeunit of *τ* now becomes important, since the timescale of a Beta(2 *- α, α*)-coalescent is proportional to *N*^*α−*1^, 1 < *α* < 2 [40], where *N* is the population size. One can obtain a more accurate expression of the timescale *given* knowledge about the mean of the potential offspring distribution [see 40]. However, since the mean is unknown in most cases, we apply the approximation *N*^*α−*1^. Assuming *n* ≥ 2 sequences from each population, the ‘observed’ FST 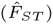 was computed as 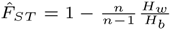 where *H*_*w*_ is the average pairwise differences within populations, 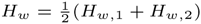, and *H*_*b*_ is the average of *n*^2^ pairwise differences between populations.

The following command-line argument for `Hybrid-Lambda` simulates 1000 genealogies with 10 lineages sampled from each of two populations separated by one coalescent unit with mutation rate *μ* = 0.00001 using a β-coalescent with parameter *α* = 1.5:

`hybrid-Lambda -spng ‘(A:10000,B:10000);’ -num 1000 -seed 45 -mu 0.00001 -S 10 10 \`

`-mm 1.5 -sim_num_mut -seg –fst`

where

- -`spng` ‘(A:10000,B:10000);’ denotes the population structure of a split model of one population splits to two at 10000 generations in the past.
- -`num` 1000 simulates 1000 genealogies from this model.
- -`seed` 45 initializes the random seed for the simulation.
- -`mu` 0.00001 specifies the mutation rate of 0.00001 per generation.
- -`S` 10 10 samples 10 individuals from each population.
- -`mm` 1.5 specifies the Λ-coalescent parameter.
- -`sim num mut` outputs simulated genealogies in Newick string, of which the number of mutations on internal branches are labelled.
- -`seg generates haplotype data set`.
- -`fst computes` *F*_*ST*_ of the generated haplotype data set.

One can use this example to generate the data for Figure 4 by setting the `-S` flag to `-S 1 1`.

